# Synchronous and Asynchronous Response in Dynamically Perturbed Proteins

**DOI:** 10.1101/2020.09.14.296111

**Authors:** Albert Erkip, Aysima Hacisuleyman, Batu Erman, Burak Erman

## Abstract

We developed a Dynamic Gaussian Network Model to study perturbation and response in proteins. The model is based on the solution of the Langevin equation in the presence of noise and perturbation. A residue is perturbed periodically with a given frequency and the response of other residues is determined in terms of a storage and loss modulus of the protein. The amount of work lost upon periodic perturbation and the residues that contribute significantly to the lost work is determined. The model shows that perturbation introduces new dynamic correlations into the system with time delayed synchronous and asynchronous components. Residues whose perturbation induces large correlations in the protein and those that do not lead to correlations may be identified. The model is used to investigate the dynamic modulation of nanobodies. Despite its simplicity, the model explains several features of perturbation and response such as the role of loops and linkers in perturbation, dispersion of work of perturbation, and information transfer through preexisting pathways, all shown to be important factors in allostery.

## Introduction

The common unifying factor in allostery is ‘perturbation and response’. When a specific site of the protein is perturbed, either statically or dynamically its effects propagate through the protein and the response of different parts of the protein to this perturbation leads to the function of the protein. Upon perturbation, new dynamic correlations emerge between the fluctuations of residue pairs. These are in addition to the existing equilibrium correlations and form the basis of allosteric response. Allostery in proteins, described as ‘The Second Secret of Life’^1^ was introduced in its earlier form by Monod, Wyman, and Changeux^2^ and Koshland, Nemethy, and Filmer^3^. In its earlier applications, allostery was conceived as a static picture where an effector molecule binds to the allosteric site, causing a conformational change at the active site where the function of the protein is performed. Since then, allostery has been a subject of continuous improvement. A major conceptual breakthrough came when Cooper et. al.,^4^ introduced dynamics into allostery after which Nussinov and collaborators focused on the dynamics of allostery, or the dynamics of perturbation and response in allostery, and illuminated several important aspects of the phenomenon with emphasis on molecular structure: Allostery without shape change^5, 6^, signal transmission and allostery^7^, the idea that all dynamic proteins are allosteric^8^, and the role of linkers and loops in allosteric transmission^9, 10^ are among the important aspects of allostery introduced and elaborated upon by Nussinov and collaborators. Experimental and computational methods for studying allosteric activity in proteins have been outlined in the review paper by Liu and Nussinov^11^. An excellent review of the original theory and its critical assessment 50 years after its introduction is given by Changeux^12^.

In the present paper, we develop a simple computational elastic net model^13^ to describe the response of proteins to dynamic perturbations which we believe will lead to a better understanding of dynamic allostery. Simple computational elastic net models have been helpful in understanding the perturbation-response mechanisms in proteins. The Linear Response^14^ and Perturbation Response Scanning^15^ models studied the response of proteins to static perturbation. Some years ago the dynamic version of the Gaussian Network Model^16^ and its frequency dependent perturbation response version^17^ was introduced. Recently, a frequency dependent perturbation response analysis model was introduced for analyzing the dynamic response to ligand binding.^18^ The present model develops these concepts where a set of residues is perturbed at a given frequency and the response of the remaining residues is predicted. The focus of the present paper is to establish a connection between dynamically (periodically) imposed perturbations and dynamic correlations that emerge as the response. The problem of allostery in this context is essentially a mechanical problem based on the solution of the Langevin equation of motion. A periodic perturbation of a residue leads to correlations between the fluctuations of residue pairs that have a synchronous component in phase with the perturbation and an asynchronous component which is out of phase.

The relationship between perturbations and work done to create the dynamic correlations plays a central role in allosteric modulation. Part of the work done by the perturbing force is recoverable due to elastic mechanisms in the protein and the remaining part is lost due to dissipative mechanisms in the system. Work done against elastic mechanisms in one cycle of loading is zero. Stated in another way, the applied force does work on the protein during the first half of the loading and the protein does work on the force during the second half of the cyclic loading, summing up to zero in one cycle. The work done against dissipative mechanisms in one cycle is nonzero and is lost or dissipated. The dissipated energy excites dissipative mechanisms and is absorbed by the protein. Some residues absorb more energy than others under the action of external forces and play important role in the function of the protein such as interacting with a ligand. The present model allows for the identification of residues that play a dominant role in energy absorption. Our calculations show that perturbations applied to loop residues are more strongly absorbed by proteins, the importance of which has earlier been shown by Nussinov and collaborators^10^. Partitioning the work of an external force into elastic and dissipative components in proteins is essential for understanding perturbation and response mechanisms in proteins but has not been studied widely. Thermodynamically, the work done by a perturbing force at constant volume goes into changing the energy and the entropy of the protein which may symbolically be expressed as:

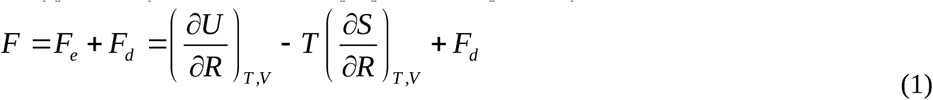

Here, *F* is the force acting on the system, *F*_*e*_ is the part of the force that goes into elastic work, *F*_*d*_ is the part that goes into dissipated work, *U* is the energy of the system, *S* is the entropy, *T* is the absolute temperature, *R* is the position variable. The first two terms on the right hand side are the elastic work components which lead to zero work in one cycle of the force. The last term is the force component corresponding to dissipated work absorbed by the protein. The first term on the right hand side of Eq. 1 is the component resulting from changes in bond lengths, bond angles bond torsion angles, and interatomic distances and contributes to the energetic part of the elastic work. The second term on the right hand side of Eq. 1 is the part resulting from changes in the probability of conformations without energy changes is the entropic elasticity part. Recently, entropy as a function of probability of conformations has been related to allosteric communication in proteins.^19, 20^ The third term on the right hand side, *F*_*d*_ is the force that excites dissipative mechanisms in the protein and leads to work lost.

A cyclic force generates time dependent correlations in the system. These correlations are in addition to the equilibrium correlations and vanish in the absence of perturbation. The generated correlations may be divided into two classes, (i) primary, and (ii) secondary. Primary correlations are between the perturbed residue and others. Secondary correlations are between two residues not directly perturbed by the force. The time dependent correlations will have a synchronous and an asynchronous component with respect to the applied force. Synchronous correlations will be in phase with the applied force, the asynchronous ones being out of phase. Asynchronous correlations vanish in the absence of dissipative mechanisms. The synchronous component contains both elastic and dissipative components, the elastic component dominating at lower frequencies of the applied force, as will be discussed in detail in the example below. Our model shows that perturbation at any site on the protein does not create new pathways but only shifts the pre-existing correlations and supports the hypothesis of pre-existing pathways proposed by Nussinov and collaborators.^9, 21^

We apply the model to a class of proteins known as nanobodies. Nanobodies are small (15kDa) proteins derived from single chain antibodies produced by members of the Camelidae family (*Llama glama, Vicugna pacos, Camelus dromedaries, Camelidae, Camelus bactrianus*), nurse sharks, *Ginglymostoma cirratum*, wobbegongs, *Orectolobus* and spotted ratfish, *Hydrolagus colliei*. ^22^ They are highly specific and have high affinity towards their target. They can be easily expressed in microorganisms for biochemical purification or intracellularly in target cells. Their toxicity is low, and their tissue penetration is not limited due their small size. Due to their biochemical functionality and economic benefits, interest in nanobodies has grown in biotechnology and medicine. In nature, binding of nanobodies to their targets takes place through their three loops known as the ‘Complementarity Determining Regions’, (CDR’s). These three CDR’s are the known regions of interaction with target molecules.

Optimization of nanobody binding can be achieved by mutating CDR residues to make the protein more suitable (i e. have higher affinity) for binding its target. These modifications which affect the three CDR loops could act as a perturbation on the protein and potentially lead to changes in the conformations and dynamics of the remaining parts of the nanobody. In the examples below, we focus on the CDR loops of nanobodies. In the simple examples we study here, we try to answer the question ‘What inter-residue correlations are introduced into the system upon external dynamic perturbation and what is their functional relevance?’ Thus, the specific aim of the present paper is to predict the perturbation-response behavior of nanobodies and changes in their dynamics. The simple structure of nanobodies allows us to trace how information flows from the energy absorbing loops to the rest of the protein. A nanobody is too small to be classified as a true allosteric protein, which is usually a large oligomeric structure of several domains as outlined by Changeux^12^. Nevertheless Tsai, Sol and Nussinov showed that all dynamic proteins are potentially allosteric^7^, and we assume the nanobody protein is allosteric in this sense. The examples we study here are limited, yet they hint to the identification of universal dynamic-response features in a given family of proteins. Although the main interest in nanobodies is on their interactions with target proteins, in this paper we focus only on the perturbation-response of nanobodies but not on their binding to their targets. We studied the latter in a separate paper where we developed a general molecular dynamics technique of predicting binding of nanobodies to their targets^23^.

## Methods

### Proteins studied

We performed calculations on several nanobodies and report results for three representative examples. Their protein Data Bank codes are, 4I0C.pdb, 5O0W.pdb, and 4KRO.pdb. 4I0C.pdb is from Arabian camel source^24^ and binds to amyloidogenic regions of the protein human lysozyme and inhibits amyloid fibril formation. 5O0W.pdb is from Alpaca and is effective in curing disease arising from the action of triponosoma^25^. 4KRO.pdb is from Llama and binds to the extracellular domain of human EGFR^26^. Their three dimensional structures are similar as shown in Figure 1 for 4I0C

**Figure 1.**
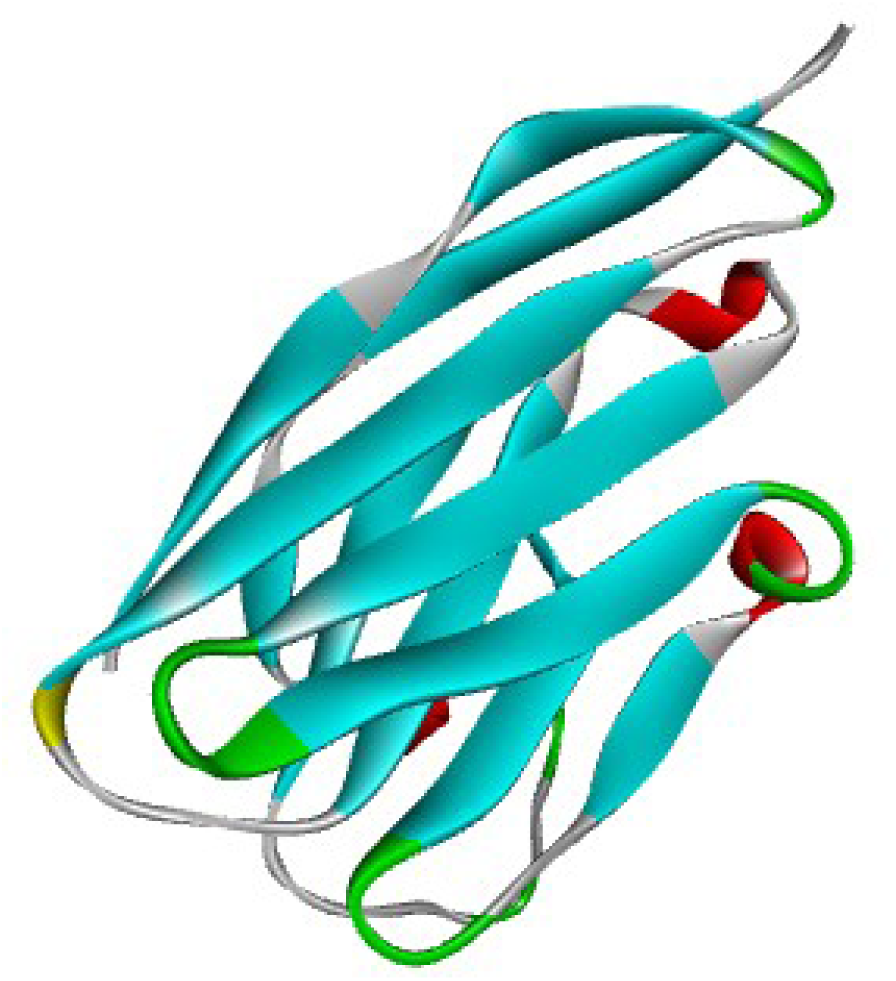
Ribbon diagram of the nanobody 4I0C.pdb. Loops are shown by green, beta strands by blue and helical structures by red. The three loops at the lower part of the figure are the three Complementarity Determining Regions (CDR’s) that bind to target proteins. The structure of other nanobodies are similar, with major difference in the length of the third CDR loop.

#### Computational tools

### The Gaussian Network model

The GNM ^13^ assumes that amino acid residues are connected to their neighbors with linear elastic springs and each residue fluctuates in space under the action of these springs. It is based on deriving an inter-residue interaction matrix or the Kirchoff matrix, by determining the number of spatial neighbors of a given residue that lie within a sphere of a given distance. It is a coarse-grained technique where all residues are collapsed on their alpha carbons. This calculation is repeated for all of the N residues. The distance, referred to as the cutoff radius, *r*_*C*_, is generally taken between 7.0 and 7.2 Å.^13^ The latter is chosen here.^27^ The connectivity or the Kirchoff matrix, Г, is obtained from the calculated distances as:

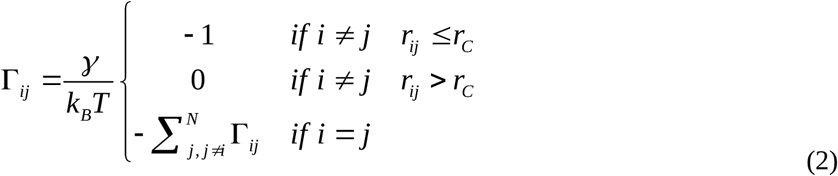

where *r*_*ij*_ is the distance between residues i and j, *γ* is a proportionality constant referred to as the spring constant of a virtual spring that represents the interaction between two neighboring residues, *k*_*B*_ is the Boltzmann constant and *T* is the absolute temperature. In all calculations here, we take 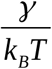 as unity because its actual value is immaterial for the purpose of this paper. In the simplest application of the GNM, we assume complete isotropy of fluctuations, with no spatial preferences. More realistic anisotropic models are present in the literature ^28^ but are not directly relevant for our model at this stage.

### Dynamics of the Gaussian Network Model of proteins

The motions of the residues of a protein obey the equation

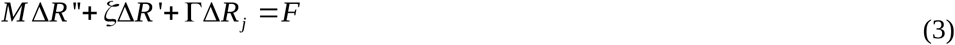

where, the prime is the time derivative, *ζ* is the friction coefficient, and *F* is the external force acting on the i^th^ residue. In an atomistic picture of the model, the force will be applied to an atom. In the present coarse-grained treatment, atoms are replaced by residues, each residue being represented by its alpha carbon, N in number. Although Δ*R*(*t*)is an 3Nx1 vector corresponding to the X, Y, and Z coordinates, as Г is isotropic, it is possible to consider Δ*R*(*t*) as the Nx1 vector corresponding to the X (and respectively Y and Z) coordinates. *M* is a diagonal NxN matrix, whose entries equate to the masses of the residues. *ζ* is the friction force acting on each residue. Since the masses are several orders of magnitude smaller than friction forces,^29^ the first term is omitted and the equation of motion, i.e., the Langevin equation, reads as

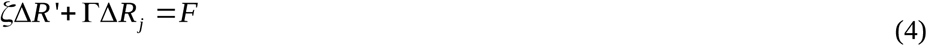

The dimensions of *ζ* are (force)(time)/(length) and the dimensions of Г are (force)/(length),, i.e., the spring constant.We consider a particular force of the form *F* = *F*_*p*_ cos (*ωt*)acting on the p’th residue with *F*_*p*_ denoting the amplitude of the force on residue p.

The solution of this equation leads to the fluctuation, Δ*R*(*t*), of residues as

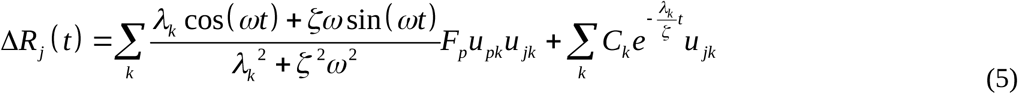

Here, Δ*R*_*j*_(*t*) is the x (and y,z) coordinate of the displacement of the jth residue, *λ*_*k*_ and *u*_*jk*_ denote the eigenvalues and the jth component of the normalized eigenvector u_*k*_. The derivation for the expression for Δ*R*(*t*)is given in Reference ^17^. The second sum in Eq. 5 decays exponentially in time and thus will have no effect in the time averages that we will compute later. For that reason, we ignore this second term.

### Work of perturbation

Eq. 5 may be written as

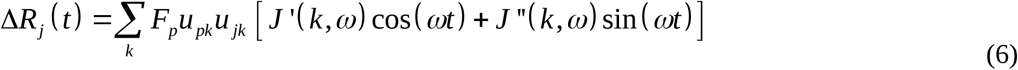

Where,

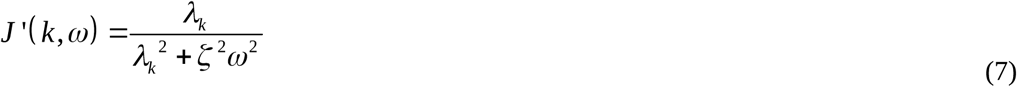

and

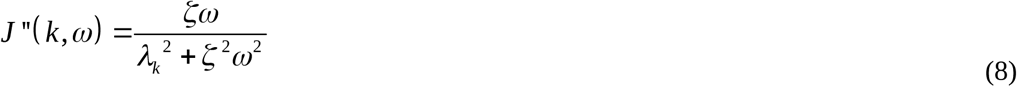

are the storage compliance and loss compliance, respectively. In both *J’* and *J’’*, the product *ζ ω* appears which has dimensions of (force)/(length), the same as that of the spring constant of the GNM. We now want to evaluate the work done defined as

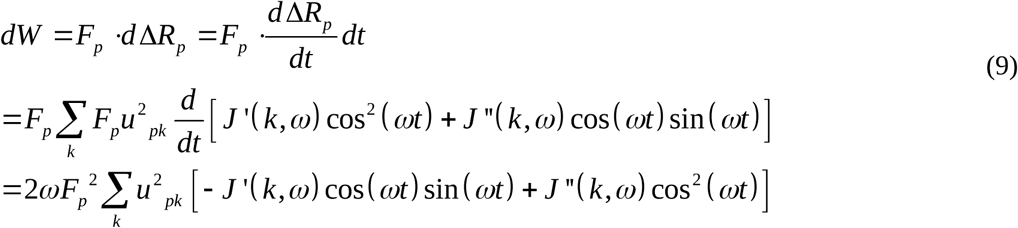

In one cycle, the work done is

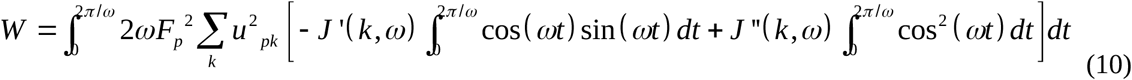

Since the first integral (energy stored in one cycle) on the right hand side of this equation is zero, the dissipated energy is obtained by the second term. The dissipated energy is then

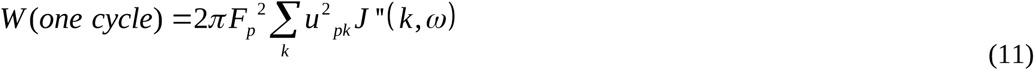

To emphasize the importance of dissipation in allostery in more detail, the dissipated work is that part of the work of the force acting on residue p that is dissipated as heat to the surroundings. Assume a ligand bound on residue p and exerting a force *F*_*p*_ on the residue. Assuming that this force is periodic, and its work on the residue p is fully dissipated, there will be no elastic response from the protein, and the ligand will not receive any feedback from the protein. The model we present can identify the pathways through which the effect of the force diffuses into the protein as will be shown in the example below.

### Synchronicity and Asynchronicity

Application of a time dependent perturbation on the protein generates new correlations among residue pairs. These correlations are in excess of the already existing static correlations in the system. Static or equilibrium correlations are expressed as^30^

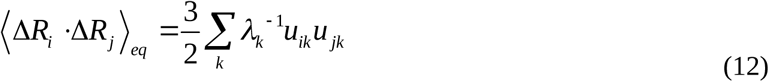

Where the angular brackets denote averaging over all equilibrium conformations of the protein. The newly generated correlations in excess of ⟨Δ*R*_*i*_ Δ*R*_*j*_⟩_*eq*_ depend on the type of the perturbation. If the time dependent perturbation is applied and removed, the correlations will be transient and decay to zero. If the system is perturbed continuously, as in the case of a cosine wave, then the new correlations will persist and will be a function of time, dependent on the perturbation. If the correlation of the fluctuations of two residues are observed at a time lag of *τ*, then the time delayed correlation of residue fluctuations will be expressed as:

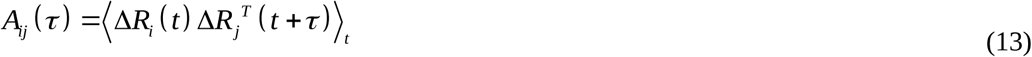

Here, the angular brackets denote average over the full trajectory of motion, the subscript t indicating that the average is taken over all time, t. The superposed T represents the transpose of the vector and *A*_*ij*_ (*τ*) is the time delayed correlation of fluctuations between residues i and j where the j ^th^ residue is observed a time *τ* after the ith residue. Substituting the displacement given by Eq 6 into the correlation expression, Eq. 12, we obtain

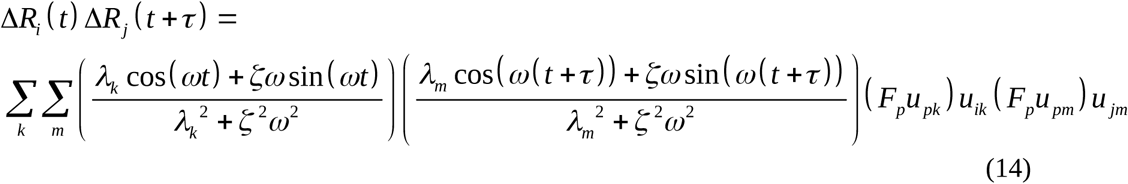

The term *u*_*ik*_ is the i^th^ element of the kth eigenvector, with similar definitions for the others. Averaging Eq. 14 over time, t, leads to

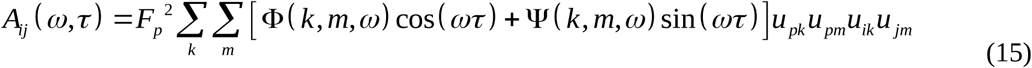

where, Φ(*k,m, ω*) follows the external perturbation and Ψ(*k, m,ω*) is out of phase with it.In Eq. 15, 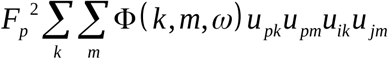 is defined as the synchronous and 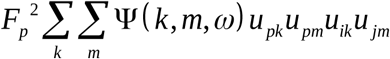 the asynchronous components of time dependent correlations, where:

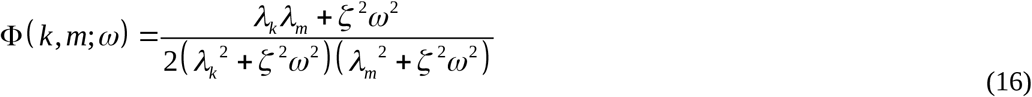

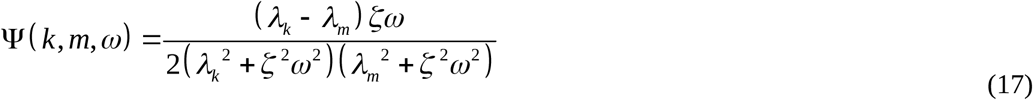

Using the definition for *J* ′(*k,ω*) and *J* ″(*k,ω*) given by Eqs 7 and 8, we obtain

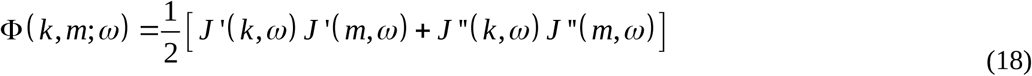

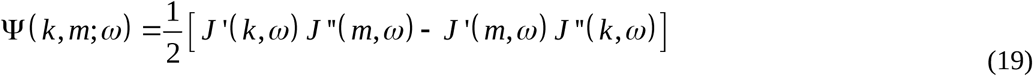

Some properties of time dependent correlations *A*_*ij*_ (*ω, τ*) are as follows: (i) the magnitude of correlations depend on the amount of time between two observations, (ii) they vanish in the absence of perturbation, (iii) they converge to 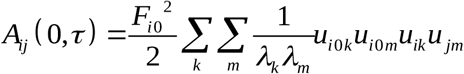 as the frequency of perturbation goes to zero. This limit may be regarded as correlations originating under static loading, (iii) the synchronous component of 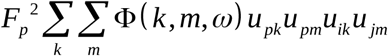, shows the correlations that are in phase with perturbation. They are symmetric in i and j and may be positive or negative, meaning that the instantaneous fluctuations of i and j follow each other or are in opposite directions, (iv) the asynchronous component, i.e., 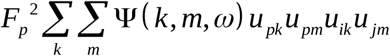 shows the correlations that are out of phase with the applied perturbation. As may be seen from Eq. 19, they are asymmetric in i and j, i.e., *A*_*ij,asynchronous*_ = − *A*_*ji,asynchronous*_. Physically, this renders a directionality to pair correlations depending which residue we observe first. This imposes a causality to correlations, i.e., there is a difference in the sign of correlations if i is observed first then j or if j is observed first and then i. This is one of the halmarks of asynchronicity. Relationship of this directionality to the causality observed in information or entropy transfer in proteins is apparent.^19, 20^ Finally, the most important feature of asynchronous correlations is that they vanish in the absence of dissipation. Therefore, asynchronicity is purely a result of dissipative mechanisms in the system. Similarly, asynchronous response vanishes when the frequency goes to zero.

### Energy absorbing pair correlations

The asynchronous component of correlations is proportional to the work dissipated and gives information on the residue pairs of residues whose interaction dissipates the work applied to the system. In order to estimate the pair interactions that lead to dissipation, we define the asynchronicity ratio as

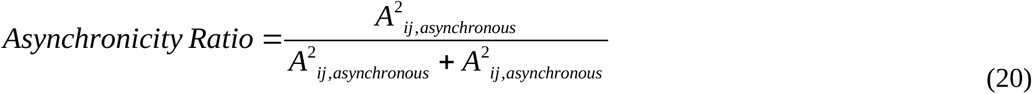

This ratio indicates the fraction of asynchronicity in the correlations. Since the asynchronous component is proportional to *J* ″(*ω*), it may be regarded as dissipative correlations.

## Results

Most of the results reported in this section are for the nanobody 4I0C and for residue 26. Residue 26 is a Glycine at the beginning of the first CDR loop and plays a role in the mechanical behavior of the protein. Results for other proteins and perturbation of different residues are qualitatively similar, but differ quantitatively. For space reasons, we focus mostly on 4I0C and on residue 26, with consderation of others when needed.

### Loops dissipate the largest amount of the energy of perturbation

In Figure 2 the amount of work dissipated in one cycle is presented as a function of residue index for the three nanobodies, 4i0c, 5o0w, and 4kro. Calculations are performed according to Eq. 11 for a friction coefficient of *ζ* =1and a frequency of *ω* =1. Each residue is perturbed, one by one, with a cosine wave cos(*ω*t) of frequency *ω*. Several maxima are observed in the curves denoting the residues that dissipate the highest amount of applied work. Those residues are in the loop regions of the nanobodies.

**Figure 2.**
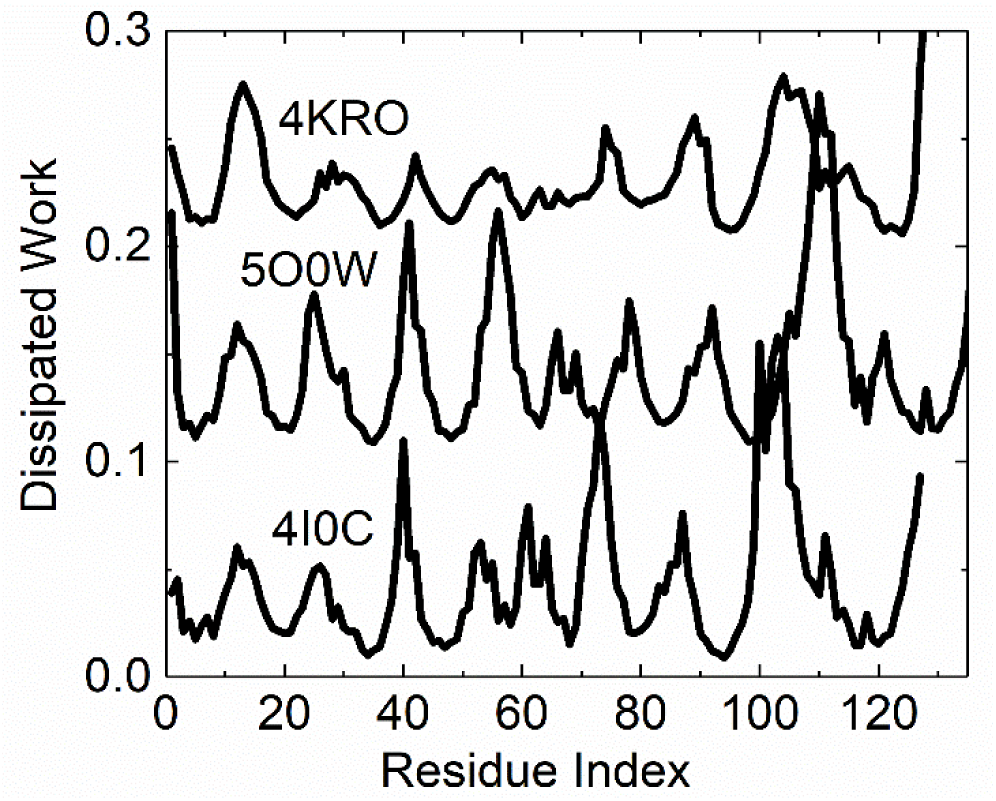
Dissipated work in one cycle.Eq. 11 is used in obtaining the curves with F_p_ = 1, *<u>ω</u>*=1 and *ζ*=1. The curves are shifted up arbitararily for clarity.

The amount of dissipation depends on the frequency of the applied perturbation as acknowledged by Eq. 8. In Figure 3 the amount of dissipated work is presented as a function of perturbation frequency obtained by Eq. 11. We analyzed results for perturbation of all residues. Here, we show the results for residue 26. At zero frequency, which corresponds to static perturbation, dissipated work vanishes. Dissipation also goes to zero, slowly, at high frequencies.

**Figure 3.**
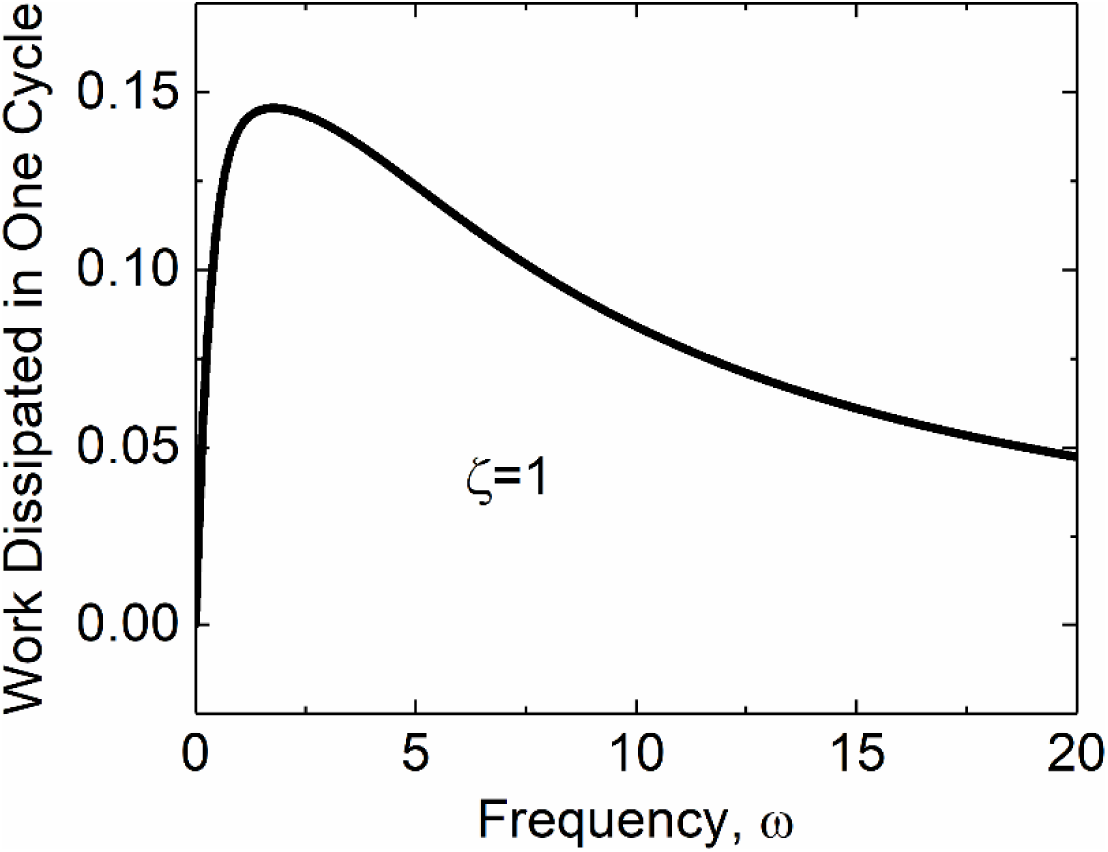
Energy dissipated in one cycle for 4I0C.pdb as a function of frequency. Eq. 8 is used in obtaining the dissipation-frequency curve.

### Periodic perturbation of a residue introduces correlations

The synchronous and asynchronous components of the pair correlations that are generated by periodically perturbing a residue are presented in Figures 4 and 5. These surfaces are obtained from Eq. 15 for p=26, *<u>ω</u>*=1 and *ζ*=1. The indices of residue pairs are indicated along the two axes in each figure and the corresponding value on the surface indicate the magnitude of the correlation produced. Perturbation of residue 26 results in its coupling with several residues in the nanobody, the strongest of which being with the amino terminal residues and residue 98. These are the primary correlations induced directly by the perturbed residue. Residue 98, although not directly perturbed, exhibits secondary correlations with several others in the nanobody as may be seen from the several peaks on the surface along the index 98 line. We now analyze direct and indirect correlations in more detail.

**Figure 4.**
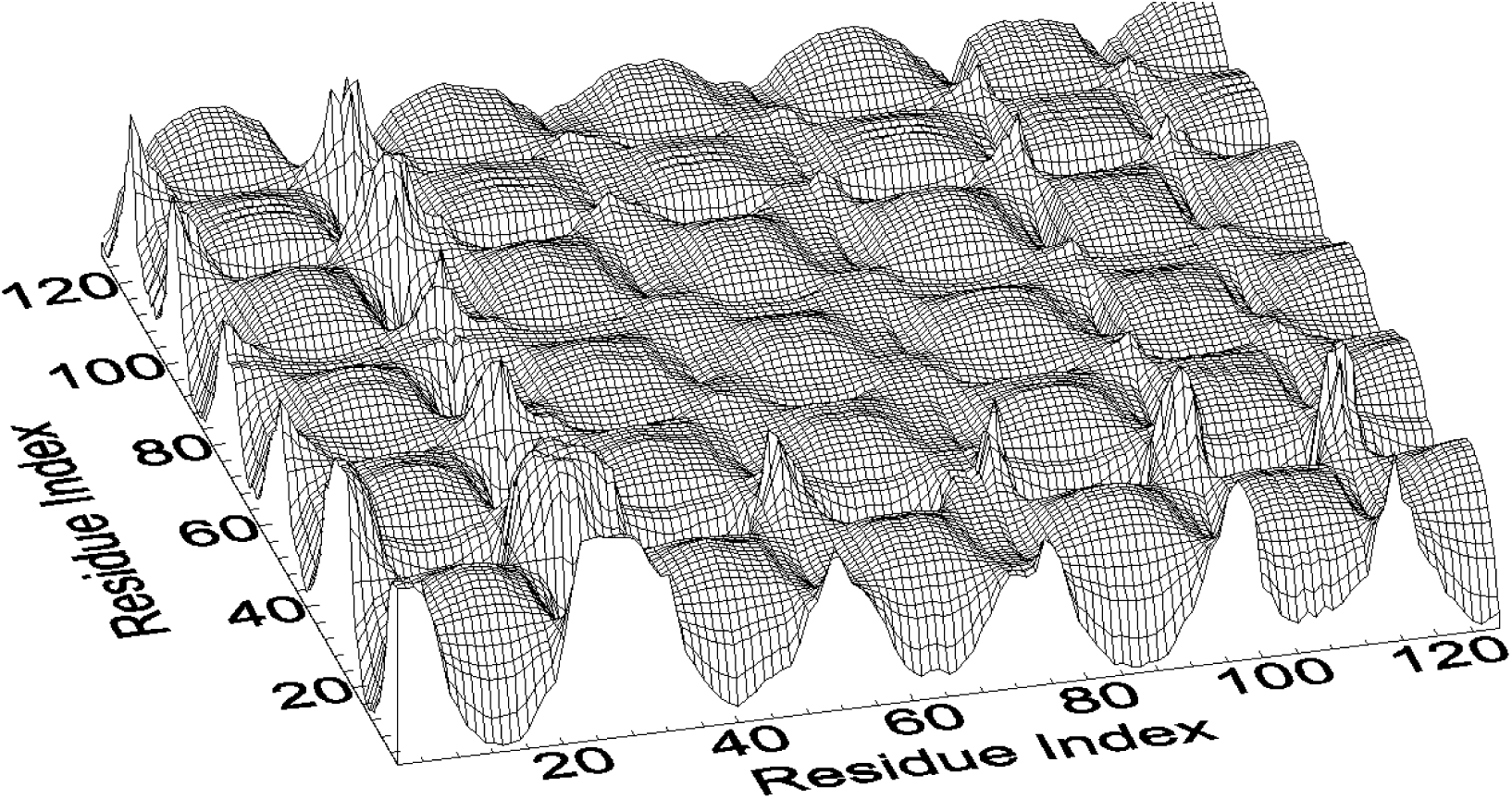
Synchronous response to dynamic perturbation of residue 26 of 4I0C.pdb.

**Figure 5.**
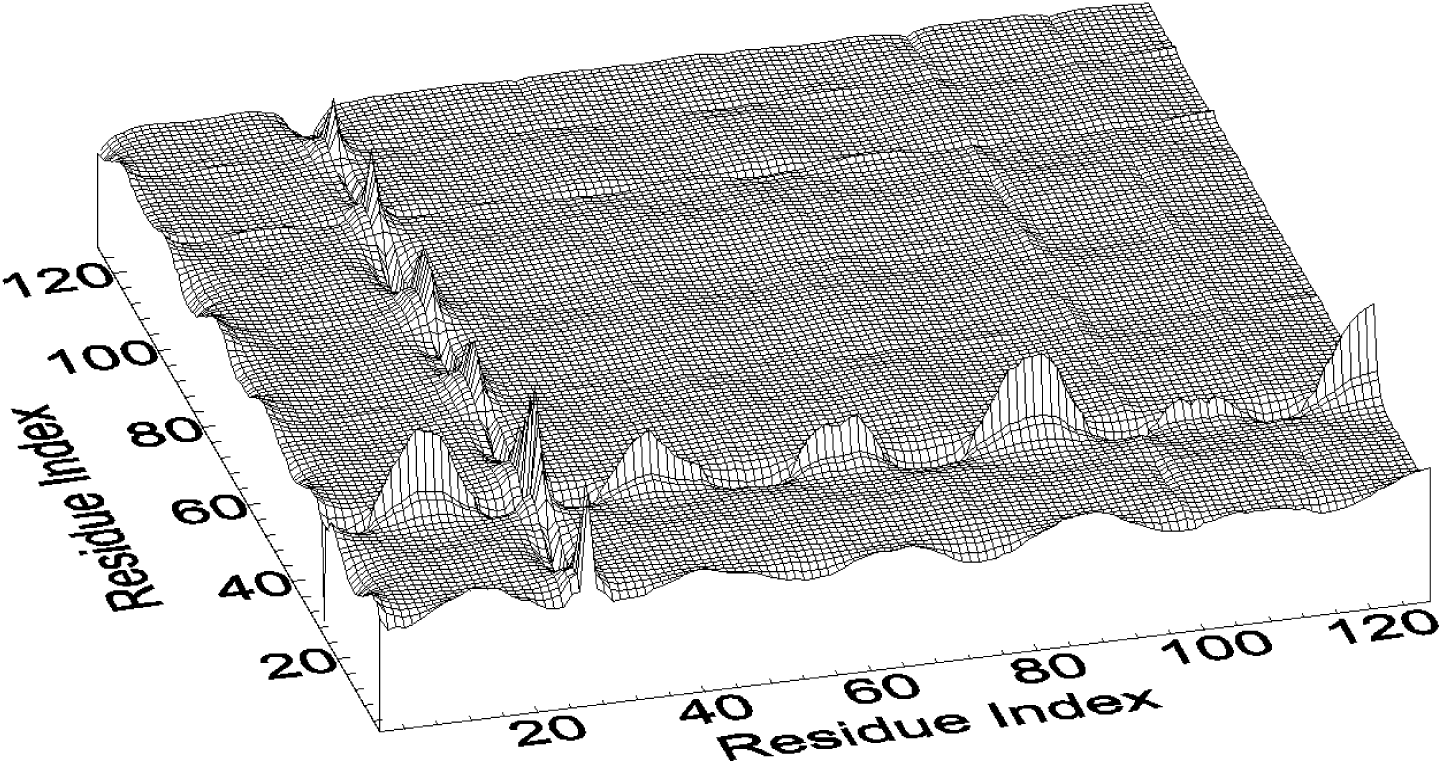
Asynchronous response to dynamic perturbation of residue 26 of 4I0C.pdb.

In figure 6 the primary correlations of residue 26 with others are presented. The synchronous and asynchronous components have their maxima and minima at the same residues. When 26 is perturbed its fluctuations are positively correlated with residues 2, 52, 73, 98 and 114 showing that they move in the same direction with 26. All of these residues are on the CDR loops of the nanobody. In addition to the positively correlated residues, 26 shows negative correlations with residues 12, 40, 61, 86, and 130. These are the residues on the non-CDR loops that are in the opposite end of the nanobody far from 26, the point of perturbation, showing that perturbation carries information to points far from the point of perturbation. An important feature of the two curves in Figure 6 is that perturbation of 26 introduces the largest correlations with CDR loop residues both for the synchronous and the asynchronous components.

**Figure 6.**
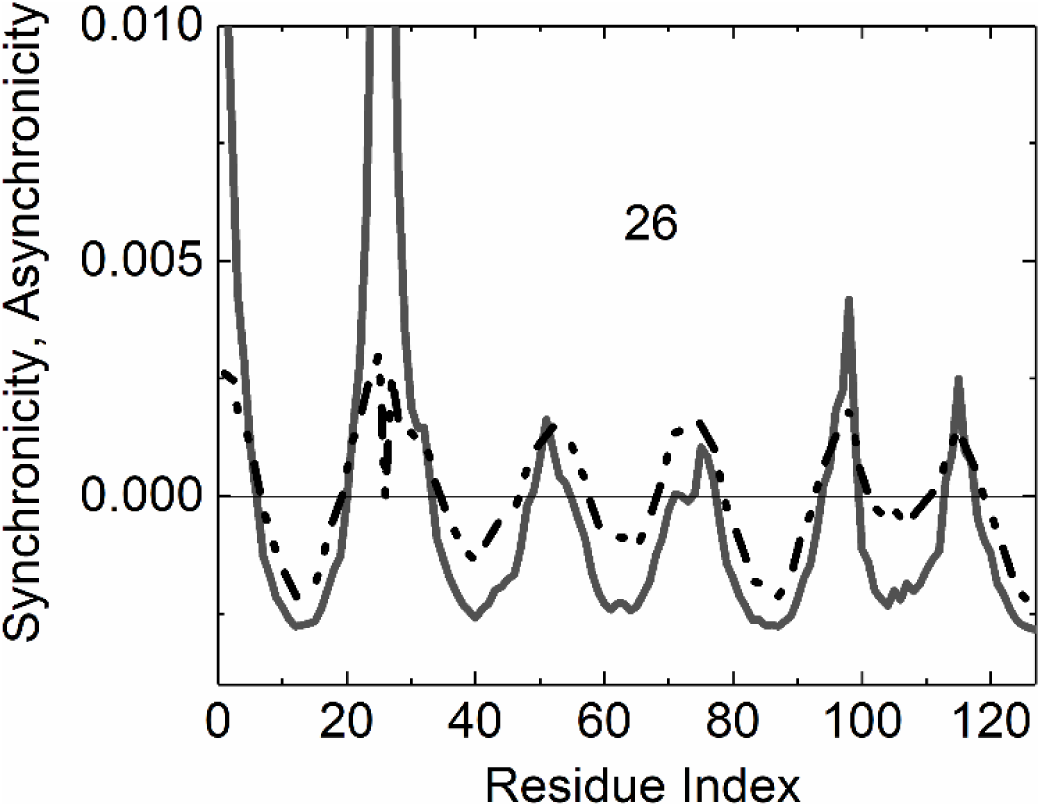
Primary correlations between residue 26 and others in the protein. The synchronous and asynchronous components are presented separately, the solid curve for the synchronous and dot-dashed curve for the asynchronous component.

In Figure 7, the secondary correlations of residue 98 with others are presented. Although residue 98 is not directly perturbed, perturbation of 26 induces correlations between different pairs, and 98 is the one that exhibits a strong secondary correlation with several residues of the protein. When 26 is perturbed, fluctuations of 98 are positively correlated with the CDR loop residues 2, 52, 73, 98 and 114 and negatively with residues 12, 40, 61, 86, and 130. The latter are the residues on the loops that are in the opposite end of the nanobody. Positions of highly correlated and anticorrelated residues are shown in Figure 8.

**Figure 7.**
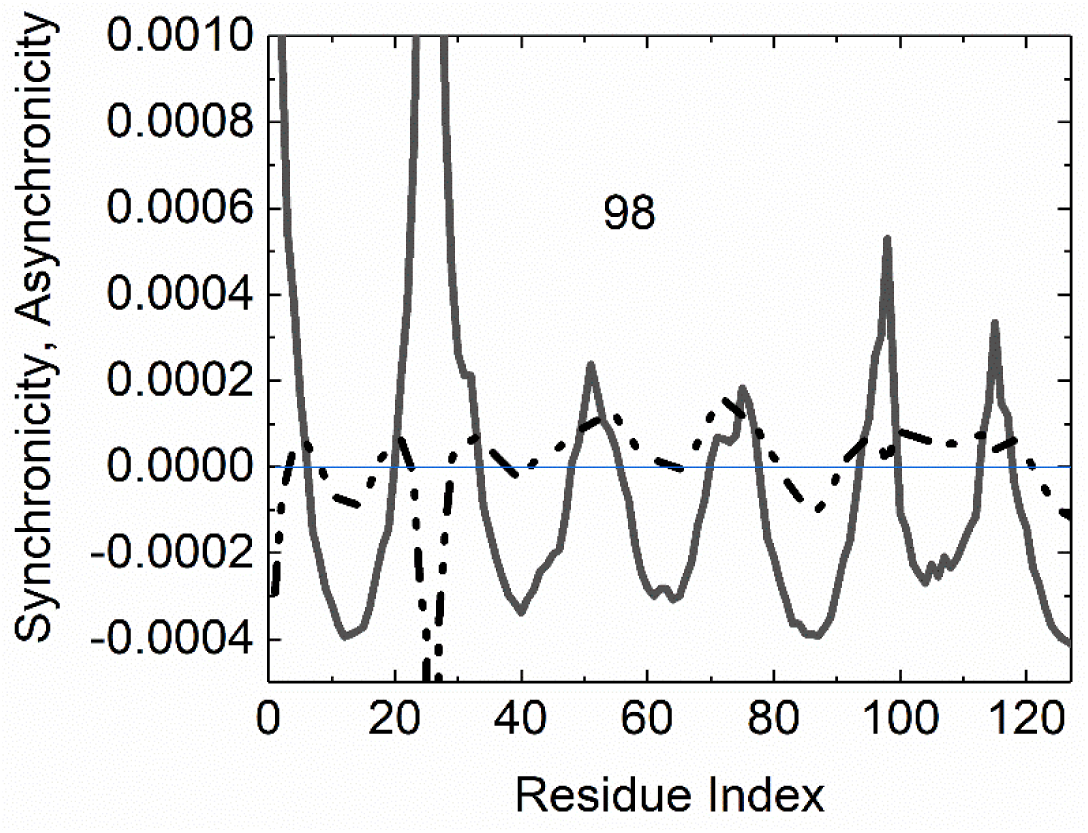
Secondary correlations between residue 98 and others in the protein upon perturbation of 26. Solid curve the synchronous, dot-dashed curve for the asynchronous component.

**Figure 8.**
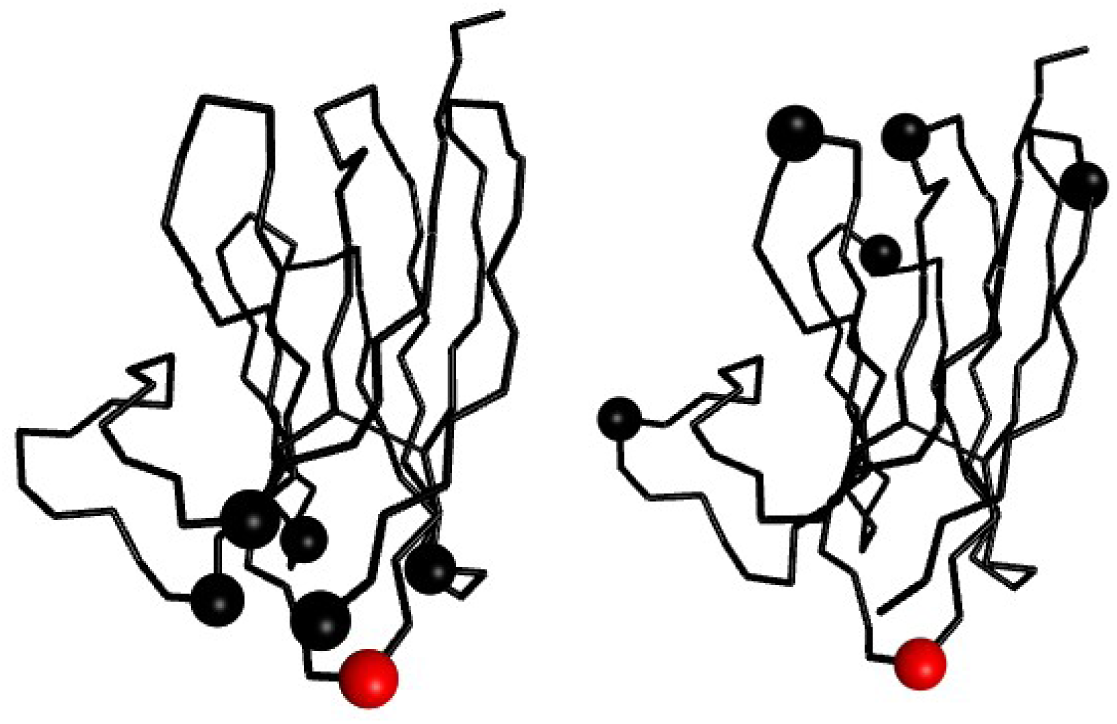
Positions of correlated (left panel) and anticorrelated (right panel) residues relative to residue 26 shown in red.

### Strong signal and no signal residues

When residues are perturbed, one by one, primary correlations are produced with other residues of the protein. The correlations produced have synchronous and asynchronous components. The latter contributes to the dispersion of the work of perturbation. For information transfer by perturbation, we now focus on the synchronous component. Some induced correlations or signals will be strong, meaning that information can be transmitted from the perturbed residue and some are weak or approximately zero indicating that there is no communication from the perturbed residue. The example for perturbation of residue 26 is elaborated in Figure 6 where the loop residues were most affected by the perturbation. Here, we consider the generated correlations from the perturbation of each residue of the protein. Figure 9 is obtained by taking the synchronous component Eq. 15:

**Figure 9.**
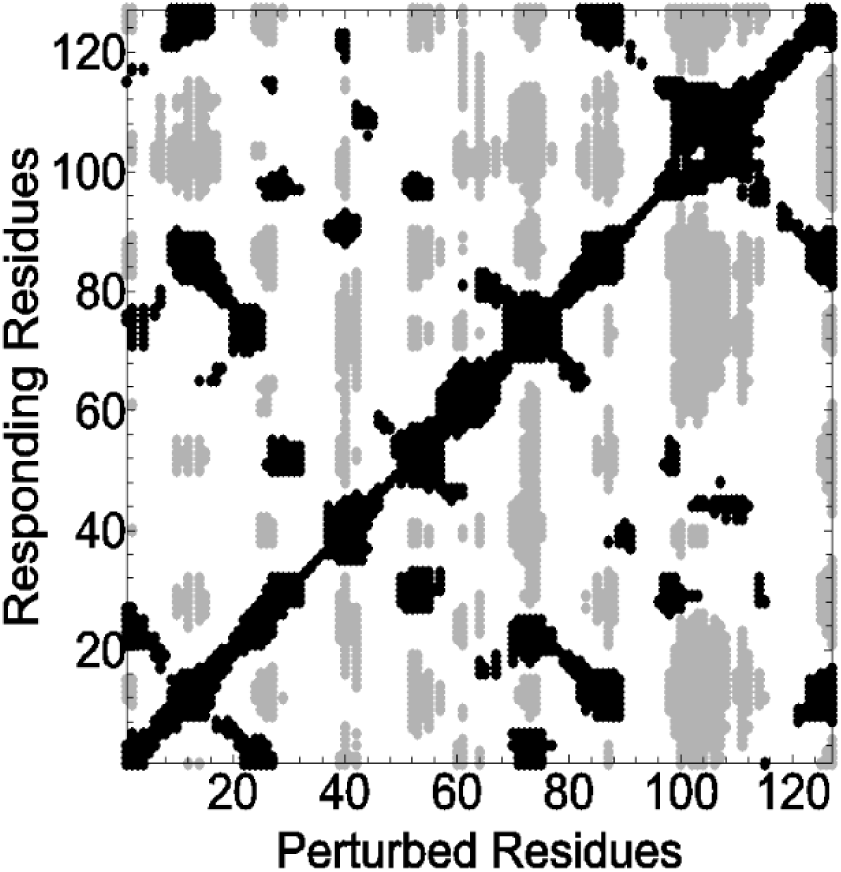
The signal and no signal residues of the nanobody obtained by Eq. 21.

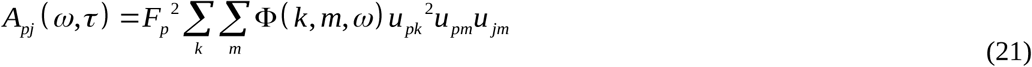

Here, a residue p is perturbed and its correlations with another residue j is calculated and the calculations are repeated for p from 1 to N. The abscissa in Figure 9 indexes the perturbed residues, the ordinate indexes the residues that show correlations with the perturbed residues. The gray dots represent negatively correlated pairs and the black dots represent positively correlated pairs. The empty regions of the graph shows the no-correlation regions. The residues whose perturbation cannot generate primary correlations are termed as no-signal residues. The points shown in Figure 9 cover the following ranges; positive correlations fall between 0.01 to 0.15, negative correlations fall between -0.15 to -0.01. We assume that correlations below ±0.01 are too small and may be regarded as negligibly small. The residues that fall in the residue ranges 29-37, 43-51, 79-82, 90-97, and 119-123 along the abscissa correspond to empty regions in the graph and these residues are therefore no signal residues. When plotted on the three dimensional pdb structure, these are observed as the residues that are located mostly on the beta strands of the nanobody.

### Asynchronous correlations with no synchronicity

The residues whose correlations with the perturbed residue dissipate most of the work of perturbation while producing little or no synchrony is of special interest. In the interest of determining these residues, we introduced the asynchronicity ratio by Eq. 20. In Figure 10 the asynchronicity ratio is presented as a function of residue index when residue 26 is perturbed. Alternatively, we locate the zeros of the synchronous curve in Figure 6. The plot shows that residues 20, 33, 49, 55, 71, 77, 94, 99, 113 and 117 are residues with zero synchronicity and nonzero asynchronicity. All of these residues are located in the beta strands that connect to the loops. We define the termini of the loops as hinge points, i.e., points where the flexible loop is anchored into the main structure of the protein.

**Figure 10.**
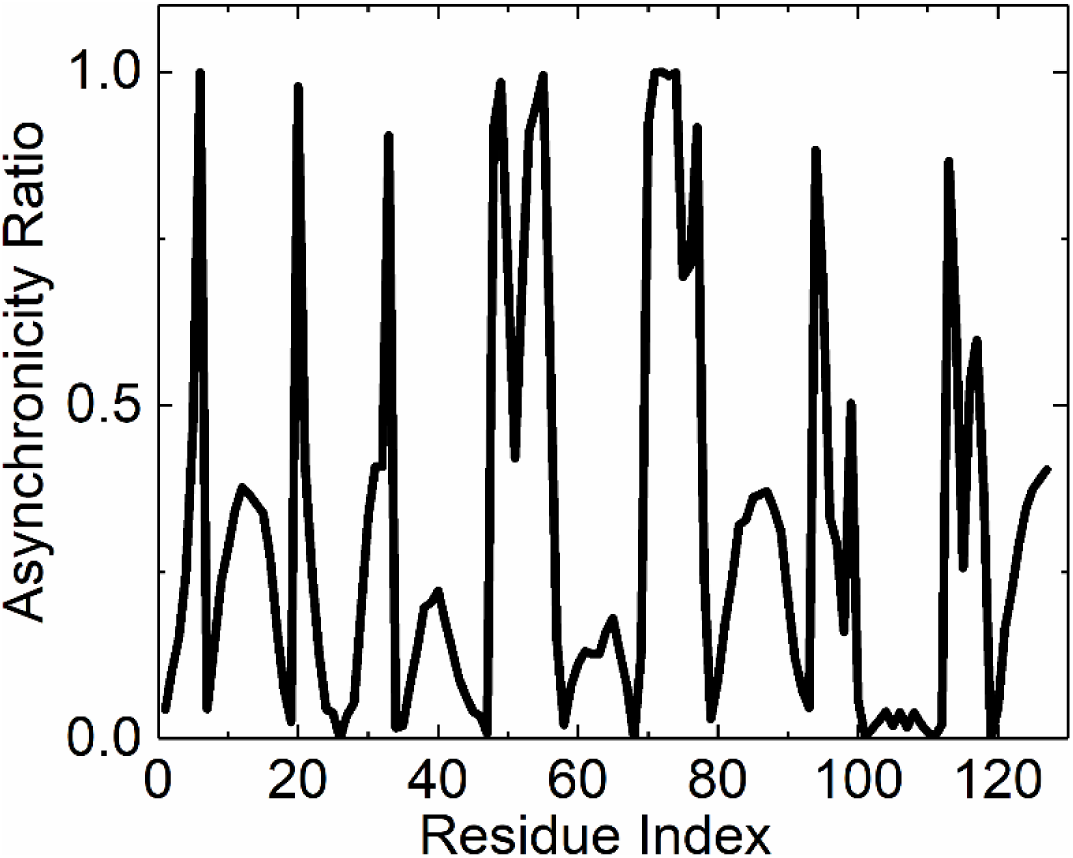
Relative asynchronicity of resides in correlations arising from the perturbation of residue 26.

### Correlations with no asynchronicity

Also of interest is the determination of residues that have no or negligible asynchronicity which means that the effects of perturbation reach those residues without any dispersion. These may be identified from minima in Figure 10. A more straightforward way to calculate this is from the asynchronicity-residue index plot by identifying the zeros of the plot. For the case of perturbing residue 26, the zeros of the curve from Figure 6 are residues 19, 34, 47, 57, 68, 78, 92, 100, 112, 119. Comparison of this set with the no synchronicity set discussed in the previous paragraph shows that the no asynchronicity residues are close to the hinge points. Therefore, hinge points emerge as of special importance in the perturbation response process.

### Dynamically generated synchronous correlations satisfy the hypothesis of pre-existing pathways

The synchronous components of the correlations generated by perturbation are proportional to the already existing equilibrium correlations as may be verified by comparing the results of equations 12 and 15 which are presented in Figure 12. The thick solid curve shows the correlation of residue 26 with other residues obtained by using Eq. 12. The thin line curve are the correlations generated when residue 26 is perturbed. The thin line curve is rescaled by multiplying by 10. The two curves are essentially identical showing that any signal generated by perturbation rides on the pre-existing correlations. This property has been suggested earlier by Nussinov and collaborators in studying the mechanisms of allostery.^9, 21^ The asynchronous components also show similarities to equilibrium correlations but they depart at residues near the perturbed residues.

**Figure 12.**
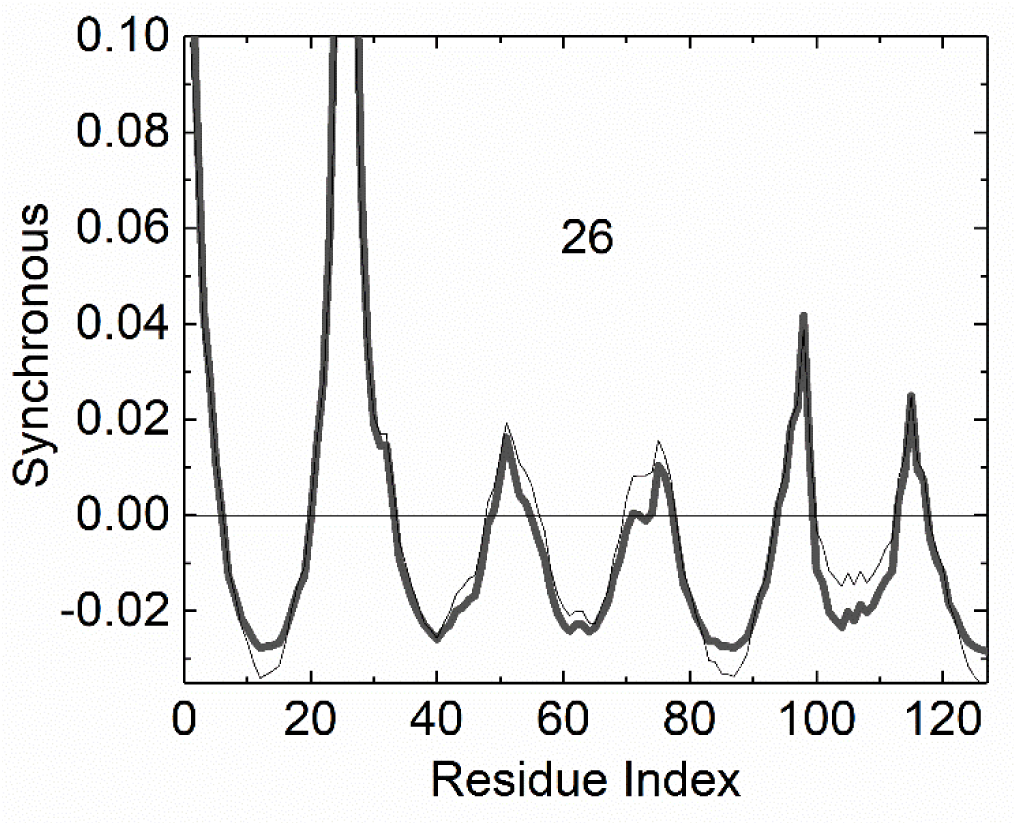
Equilibrium correlations of residue 26 with the remaining residues of the nanobody (thick line) compared with synchronous response when 26 is perturbed (thin line). The thin line curve is obtained by taking F_p_ = 1, *<u>ω</u>*=1 and *ζ*=1.

## Discussion

We used a simple isotropic elastic net model based on the GNM to predict effects of perturbing a residue in a protein. Such perturbations play an important role on signal transduction in proteins and understanding them is of primary importance. Perturbation of residues induces new pair correlations in the protein. If a pair correlation is between the perturbed residue and another one, we call it a primary correlation. If a correlation emerges between two residues upon the perturbation of another residue, we call this a secondary perturbation. Secondary perturbations are weaker than primary perturbations, but nevertheless they are nonzero and may be consequential in the function of the protein. We based our calculations mainly on the perturbation of residue 26, and for the nanobody 4I0C which are perfectly representative of all others. Therefore, we did not present the others in detail for brevity.

A time dependent perturbation induces time dependent pair correlations as shown by the model. The time dependence is best explained by synchronicity and asynchronicity of correlations with the applied perturbation. Asynchronous correlations arise from dispersive mechanisms in the system and may be used to analyse dissipative mechanisms in allosteric communication, such as dissipated work of external perturbation. The model presented in this paper points to the importance of loops in energy and signal transfer in perturbed proteins. Pair residue correlations generated by dynamically perturbing a loop residue creates a correlation of fluctuations of residue pairs, thus showing that external perturbation may be used to transmit signals to certain residues of the protein. Interestingly, some residues, those that lie on the beta strands of the nanobody cannot be used to transmit signals by perturbation. Perturbation of loops do not act locally but can create signals that reach long distances in the nanobody. This was pointed out by Nussinov and collaborators^10^ and was defined as vehicles that propagate allosteric effects.

The primary function of a nanobody is to bind to its target with high affinity. In this context, an allosteric activity for nanobodies as in the classical examples of allosteric proteins is not immediately apparent. However, the present study clearly shows that perturbation leads to signal propagation in the nanobody. As nanobodies are only the variable regions of the single chain antibodies produced in nature, perturbation of the target binding region of the single chain antibody may allosterically propagate signals to the constant region of this molecule that may have structural consequences. We showed that signals go from the CDR loops to the loops on the opposite end of the nanobody. Some years ago, Gunasekaran, Ma and Nussinov^8^ suggested that all dynamic proteins are allosteric. With the help of the model we propose herein, we would like to modify this hypothesis slightly as ‘It is possible to send dynamic signals through all proteins. These signals use the equilibrium correlations as a vehicle’.

The dynamic perturbation can be of any origin, mechanical, electromagnetic, etc. and the correlations that are generated can be analyzed by a multitude of techniques. The model introduced in this paper has several features in common with two dimensional infrared spectroscopy where an external periodic perturbation of a given frequency activates certain motions in the system the response spectrum of which can be observed by infrared correlation spectroscopy.^31^References

## References

1 J. Monod, Austryn Wainhouse (New York: Vintage) (1970)

2 J. Monod, J. Wyman, and J.-P. Changeux, J Mol Biol 12 (1965) 88.

3 D. Koshland Jr, G. Nemethy, and D. Filmer, Biochemistry 5 (1966) 365.

4 A. Cooper, and D. Dryden, European Biophysics Journal 11 (1984) 103.

5 C.-J. Tsai, A. Del Sol, and R. Nussinov, Journal of molecular biology 378 (2008) 1.

6 R. Nussinov, and C.-J. Tsai, Current opinion in structural biology 30 (2015) 17.

7 C.-J. Tsai, A. Del Sol, and R. Nussinov, Molecular Biosystems 5 (2009) 207.

8 K. Gunasekaran, B. Ma, and R. Nussinov, Proteins: Structure, Function, and Bioinformatics 57 (2004) 433.

9 B. Ma et al., Structure 19 (2011) 907.

10 E. Papaleo et al., Chemical reviews 116 (2016) 6391.

11 J. Liu, and R. Nussinov, PLoS computational biology 12 (2016) e1004966.

12 J.-P. Changeux, Annual review of biophysics 41 (2012) 103.

13 I. Bahar, A. R. Atilgan, and B. Erman, Fold Des 2 (1997) 173.

14 M. Ikeguchi et al., Physical review letters 94 (2005) 078102.

15 C. Atilgan, and A. R. Atilgan, PLoS Comput Biol 5 (2009) e1000544.

16 T. Haliloglu, I. Bahar, and B. Erman, Physical Review Letters 79 (1997) 3090.

17 A. Erkip, and B. Erman, Polymer 45 (2004) 641.

18 Y. B. Varolgunes, and A. Demir, Physical biology 15 (2018) 026009.

19 A. Hacisuleyman, and B. Erman, PLOS Computational Biology 13 (2017) e1005319.

20 A. Hacisuleyman, and B. Erman, Proteins: Structure, Function, and Bioinformatics (2017)

21 A. del Sol et al., Structure 17 (2009) 1042.

22 S. Muyldermans, Annual review of biochemistry 82 (2013) 775.

23 A. Hacisuleyman, and B. Erman, J Biol Phys (2020)

24 E. De Genst et al., The Journal of Physical Chemistry B 117 (2013) 13245.

25 J. Pinto et al., PLoS neglected tropical diseases 11 (2017) e0005932.

26 K. R. Schmitz et al., Structure 21 (2013) 1214.

27 C. Tuzmen, and B. Erman, Plos One 6 (2011) e16474.

28 A. R. Atilgan et al., Biophysical journal 80 (2001) 505.

29 M. Fixman, The Journal of Chemical Physics 42 (1965) 3831.

30 T. Haliloglu, E. Seyrek, and B. Erman, Physical Review Letters 100 (2008) 228102.

31 I. Noda, Applied spectroscopy 47 (1993) 1329.

